# Cryo-EM structure of the MinCD copolymeric filament from *Pseudomonas aeruginosa* at 3.1 Å resolution

**DOI:** 10.1101/638619

**Authors:** Andrzej Szewczak-Harris, James Wagstaff, Jan Löwe

**Affiliations:** MRC Laboratory of Molecular Biology, Cambridge, United Kingdom; Department of Biochemistry, University of Cambridge, United Kingdom

**Author notes:** Correspondence:* Jan Löwe, Structural Studies Division, MRC Laboratory of Molecular Biology, Francis Crick Avenue, Cambridge, Cambridgeshire, CB2 0QH, United Kingdom.

**Keywords:** bacterial cytoskeleton, cell division, protein filaments, WACA, cryo-EM, helical reconstruction

## Abstract

Positioning of the division site in many bacterial species relies on the MinCDE system, which prevents the cytokinetic Z-ring from assembling anywhere but the mid-cell, through an oscillatory diffusion-reaction mechanism. MinD dimers bind to membranes and, via their partner MinC, inhibit the polymerisation of cell division protein FtsZ into the Z-ring. MinC and MinD form polymeric assemblies in solution and on cell membranes. Here, we report the high-resolution cryo-EM structure of the copolymeric filaments of *Pseudomonas aeruginosa* MinCD. The filaments consist of three protofilaments made of alternating MinC and MinD dimers. The MinCD protofilaments are almost completely straight and assemble as single protofilaments on lipid membranes, which we also visualised by cryo-EM.

## Introduction

In most bacteria, cell division depends on the action of the Z-ring, which is formed from protein filaments consisting of bacterial tubulin homologue FtsZ, alongside other proteins. The Z-ring has the ability to constrict lipid membranes *in vitro*. Together with the downstream divisome complex, it also organises peptidoglycan remodelling at the division site [1–6]. To achieve symmetrical cell division, the Z-ring needs to be precisely positioned. Many bacteria contain at least one system responsible for correct Z-ring placement, with nucleoid exclusion and the Min system being most prominent [7–10].

In *Escherichia coli*, the Min system has three protein components: MinC, MinD and MinE [11,12]. The proteins oscillate between the cell poles, establishing the division site, or allowing the formation of a division site, at the point of minimum average concentration of MinC, normally mid-cell [13–16]. MinD is a deviant Walker A ATPase that dimerises upon ATP binding. Dimerised MinD can bind to the cell membrane, as its C-terminal amphipathic helix becomes activated [17,18]. MinE, which is also membrane-binding, stimulates the ATPase activity of MinD and therefore its dissociation from the membrane [19]. Released MinD will then undergo exchange of ADP for ATP and reassemble in an area devoid of MinE. Because of built-in delays and non-linearities in the system, MinDE constitute a dynamic and oscillating Turing reaction-diffusion device and lead to oscillation in confinement. The regulation of the Z-ring is effected by MinC, which acts as a direct inhibitor of FtsZ [20,21].

The precise action and MinD activation of MinC is unclear, but MinC seems to inhibit Z-ring formation by disassembling the FtsZ filaments and/or blocking their lateral interactions [21,22]. The MinC protein forms a stable homodimer and has two distinct domains: N-terminal MinC^N^ and C-terminal MinC^C^[23]. Both domains have the ability to inhibit FtsZ filamentation, albeit by different mechanisms [24]. The MinC^C^ domain forms the MinC dimerisation interface, binds the conserved C-terminal tail domain of FtsZ and, importantly, binds MinD [23,24].

MinCD filaments were first observed by Ghosal *et al*. and Conti *et al*., who showed that MinCD from *Escherichia coli* as well as the thermophilic bacterium *Aquifex aeolicus* polymerise in the presence of ATP to form a new class of alternating, copolymeric filaments, which can be assembled *in vitro* either on lipid membranes or in solution [25,26]. Similar assemblies have since been reported by Huang *et al*., in MinCD preparations from *Pseudomonas aeruginosa* [27]. These studies also indicated that MinCD co-assemble into filaments with a 1:1 stoichiometry and it was suggested that MinE has the ability to depolymerise MinCD filaments and inhibit their assembly.

Prior to the discovery of MinCD filaments, the proposed mechanism of FtsZ inhibition by the Min system involved MinD dimerisation and the resulting targeting of MinC dimers to the membrane, where they would inhibit FtsZ polymerisation. In this scenario, MinD would oscillate with MinC as a passenger, and inhibit the Z-ring formation at and near the poles, where protein concentration is highest [15,16,28]. The presence of MinCD filaments that can form on the cell membrane extends this proposition to include the possible inhibition of FtsZ filaments by activated, polymerised MinCD, especially given that the two have a matching periodic repeats of ~ 4 nm and ~ 8 nm, a fact that could cause very strong avidity and cooperativity effects [10,25,29].

The issue of MinCD filamentation and Z-ring disruption is far from resolved. In a 2015 study, Park *et al*. mutated surface residues of MinC and MinD monomers in *E. coli* that were implicated in MinCD filament formation. Their non-polymerising, dimer-asymmetric MinCD mutants still inhibited Z-ring formation, suggesting that the filamentation of MinCD may not be necessary for the activity of the system in cells, putting into question the earlier proposal of a function for MinCD filaments in the activation of MinC [30].

Thus far the evidence surrounding MinCD copolymeric filaments has come from biochemical experiments with purified components, such as filament pelleting or light-scattering assays, with structural data limited to low-resolution electron microscopy images [25,27]. A hybrid model for a MinCD filament has been proposed, based on the crystal structure of MinC in complex with MinD, but the resulting alternating MinC_2_-MinD_2_ protofilament model is strongly bent and did not fully recapitulate the observed EM images, further weakening the argument [25]. The discovery that MinCD from *P. aeruginosa* forms filamentous assemblies *in vitro* motivated us to investigate the structural basis for MinCD filament formation at high resolution with electron cryo-microscopy (cryo-EM).

For this study, we imaged MinCD filaments in solution and obtained a refined atomic model of the polymerised filament at 3.1 Å resolution. Additionally, we polymerised MinCD filaments on the surface of narrow lipid nanotubes and imaged the MinCD-decorated tubes with cryo-EM, verifying the membrane binding mode of single MinCD protofilaments.

## Materials and Methods

### Protein expression and purification

Full-length MinC and MinD from *P. aeruginosa* were cloned as described previously [27]. The protein gene was cloned into pET-15b, yielding a fusion protein with a polyhistidine tag on the N-terminus, followed by a thrombin cleavage site (MinC: MGSSHHHHHHSSGLVPRGSH-1-263; MinD: MGSSHHHHHHSSGLVPRGSH-1-271). The tag was not removed during purification, as it has been reported to have little effect on MinCD polymerisation [27]. Both MinC and MinD were prepared and handled in the same manner. Protein expression was carried out in *E. coli* strain C41(DE3) (Lucigen) in 2×TY media supplemented with 100 μg·L^−1^ ampicillin. Cell cultures were grown at 37□ with shaking, until cell density reached OD_600_ 0.6, when the temperature was reduced to 30°C and expression was induced by addition of 0.5 mM isopropyl β-d-1-thiogalactopyranoside (IPTG). Cells were harvested by centrifugation after 5 h expression. Harvested pellets were resuspended in NiA buffer (50 mM Tris·HCl, 300 mM NaCl, 2 mM TCEP, 1 mM NaN_3_, pH 7.5) and sonicated on ice. The lysate was cleared by centrifugation at 100,000 x *g* for 45 min and loaded onto a 5 mL HisTrap HP column (GE Healthcare). The column was washed extensively with NiA buffer. Bound protein was eluted with a gradient of increasing imidazole concentration. The eluate was collected in fractions and analysed for purity and composition with SDS-PAGE. Fractions containing protein were pooled and concentrated using Amicon Ultra-15 centrifugal filter unit (10-kDa molecular mass cut-off; Merck) until total protein concentration of 10 mg·mL^−1^ was reached. The concentrate was dialysed extensively against the polymerisation buffer (20 mM HEPES·Na, 100 mM CH_3_COOK, 5 mM (CH_3_COO)_2_Mg, pH 7.0). After dialysis, purified protein was flash-frozen in liquid nitrogen.

### Cryo-EM sample preparation and data collection

For the purpose of imaging the unsupported filaments, concentrated solutions of MinC and MinD were diluted with polymerisation buffer to 0.5 mg·mL^−1^ and combined in equal proportion. Filament polymerisation was induced by addition of 1 mM ATP and followed by 15 mins incubation at room temperature. Three microliters of polymerised sample were applied onto R 2/2 holey carbon support film on a 300-mesh copper EM sample grid (Quantifoil Micro Tools), which had been glow discharged immediately prior to use. The sample on the grid was blotted and then vitrified in liquid ethane at - 180□ by plunge-freezing using a Vitrobot Mark IV (Thermo Fisher Scientific).

Grids used to image the filament on lipid nanotubes were prepared as above, however before the addition of ATP, the MinCD mixture was combined with nanotubes. To prepare the nanotube solution, *E. coli* total lipid extract (Avanti Polar Lipids) was mixed with d-galactosyl-ß-1,1’ *N*-nervonoyl-d-erythro-sphingosine (galactosylceramide; Avanti Polar Lipids) at 3:7 weight ratio and dissolved in chloroform to a total concentration of 1 mg·mL^−1^. A 100-μL aliquot of the lipid solution in a glass vial was dried carefully under a stream of nitrogen until all the solvent evaporated and the deposit in the vial was suspended in 100 μL of polymerisation buffer. The mixture was placed in a rotary mixer for 15 min at 4□ and then transferred to a sonication bath for 1 min. Before polymerisation, freshly prepared nanotube solution was mixed in equal proportion with the MinCD solution as above and incubated for 5 min on ice.

Cryo-EM grids used for MinCD structure solution were imaged with a Titan Krios G3 transmission electron microscope (TEM; Thermo Fisher Scientific) operated at 300-kV accelerating voltage and liquid nitrogen temperature. The images were recorded using automated acquisition software with a Falcon 3EC direct electron detector (Thermo Fisher Scientific) operated in counting mode. Each exposure lasted 60 s and was collected as a 50-frame movie using total electron fluence of 50 e^−^·Å^−2^ at 0.824-Å pixel size and nominal defocus range between −1.2 and −2.5 μm.

Cryo-EM grids of MinCD filaments on lipid nanotubes were imaged with a Tecnai Polara G2 TEM (Thermo Fisher Scientific) at 300-kV accelerating voltage and liquid nitrogen temperature. The images were recorded with a prototype Falcon 3 direct electron detector (Thermo Fisher Scientific) operated in linear integration mode. The exposures lasted 1.5 s and were collected as 46-frame movies with a total electron fluence of 38 e^−^ ·Å^−2^ at 1.34-Å pixel size and nominal defocus range between −1.2 and −2.5 μm.

### Cryo-EM data processing and structure solution

Filament structure solution was carried out in RELION 3.0 [31] using the helical reconstruction method [32]. In total, a dataset of 4,045 movies was collected. The images were corrected for beam-induced motion with dose-weighting inside RELION and their contrast transfer functions were estimated with Gctf on non-dose-weighted movie averages [33]. From this dataset, 3,050 images with Gctf-estimated resolution below 4.0 Å were selected. About two thousand helical segments were selected manually and classified in 2D as single particles to provide references for automated particle picking. Automatically picked helical segments were extensively 2D-classified yielding 161,543 particles, which were extracted in 480-pixel boxes. These were used for the first round of 3D auto-refinement using a simulated helix as an initial model, which was created with the *relion_helix_toolbox* utility of RELION 3.0 and low-pass-filtered to 30 Å. The first experimental 3D map revealed presence of a 2-fold symmetry axis perpendicular to the main axis of the filament, so ‘*D*_1_’ symmetry (*C*_2_ along X axis in RELION nomenclature) was applied in subsequent refinement rounds. A focused 3D classification with a mask around the filament allowed us to further improve the homogeneity of the dataset, narrowing the particle number down to 118,659. For the later rounds of refinement, we polished the particles using Bayesian polishing function in RELION 3.0 [34]. Additionally, the initial CTFs were corrected using the CTF refinement option, including per-particle defocus estimation, per-micrograph astigmatism as well as beam-tilt estimation [35]. In the final step of 3D auto-refinement a solvent mask covering the central 30% of the helix *z*-length was used to calculate solvent-flattened Fourier shell correlation (FSC) curves. The map was postprocessed with a mask covering 20% of the helix *z*-length. The overall as well as the local resolution was assessed using the gold-standard FSC procedure in RELION, using the FSC_0.143_ criterion [36].

The images of MinCD filaments on lipid nanotubes were processed in RELION 3.0, in a manner similar to the filaments in solution. The images were corrected for beam-induced motion and their CTFs were estimated as above. Manually picked particles were extracted and 2D-classified as non-helical objects.

To build an atomic model of *P. aeruginosa* MinCD filament, homology models of MinC and MinD were created using SWISS-MODEL [37]. These were then fitted into the central portion of the final postprocessed cryo-EM map as a MinD_2_-MinC_2_ heterotetramer. The surrounding part of the map was cut out using REFMAC [38]. The homology model was adjusted manually using MAIN [39] and refined in both reciprocal and real space with REFMAC and PHENIX [40]. For reciprocal space refinement, the prepared segment of the cryo-EM map was back-transformed into structure factors using REFMAC in SFCALC mode. After fitting and refining, the model was subjected to the standard *R*-factor analysis and its agreement with standard geometry was assessed with MolProbity [41]. For data collection, image processing and atomic refinement statistics see Table 1. The refined atomic coordinates were deposited in the Protein Data Bank (PDB) with accession number 6RIQ and the corresponding final EM density in the EMDB with accession number EMD-4897.

**Table T1.**
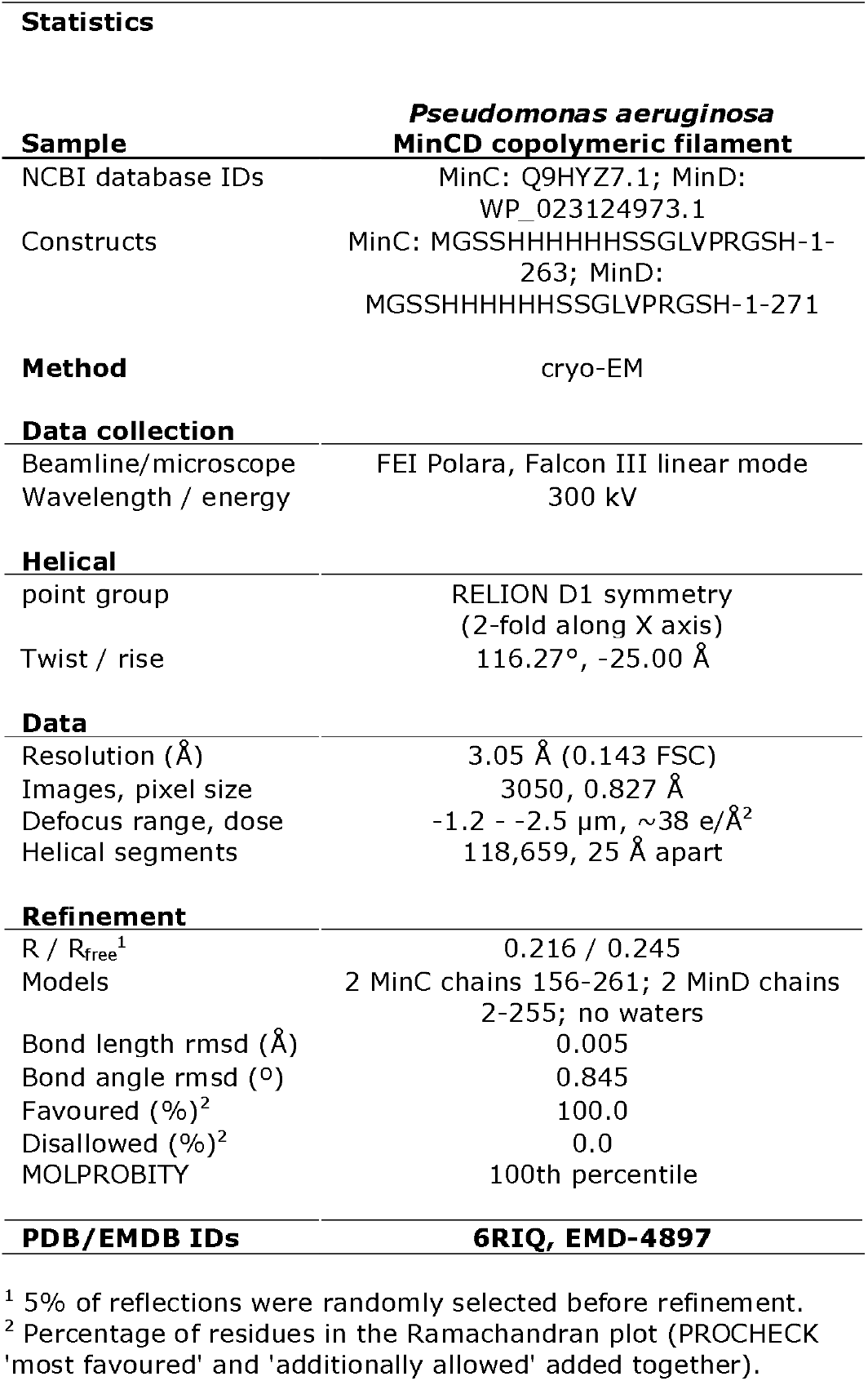
Cryo-EM and model data

## Results and Discussion

### MinCD forms triple helical filaments in solution

A previous study on MinCD copolymers from *P. aeruginosa* showed that the two proteins polymerise upon addition of ATP into a mixture of double, helically-twisted filaments, as well as seemingly single, thin filaments, visible in negatively stained electron micrographs (Fig. 1A, cf. Fig. 1A in [27]). Prompted by this observation, we set about to obtain a high-resolution reconstruction of the filaments, planning to utilise the helical nature of the filaments for their reconstruction from 2D transmission images, which has recently shown success in providing well-defined atomic structures of other helical filaments [42–45]. We purified full-length, His_6_-tagged MinC and MinD from *P. aeruginosa* and imaged their ATP-polymerised assemblies using EM (Fig. 1B).

**Fig. 1.**
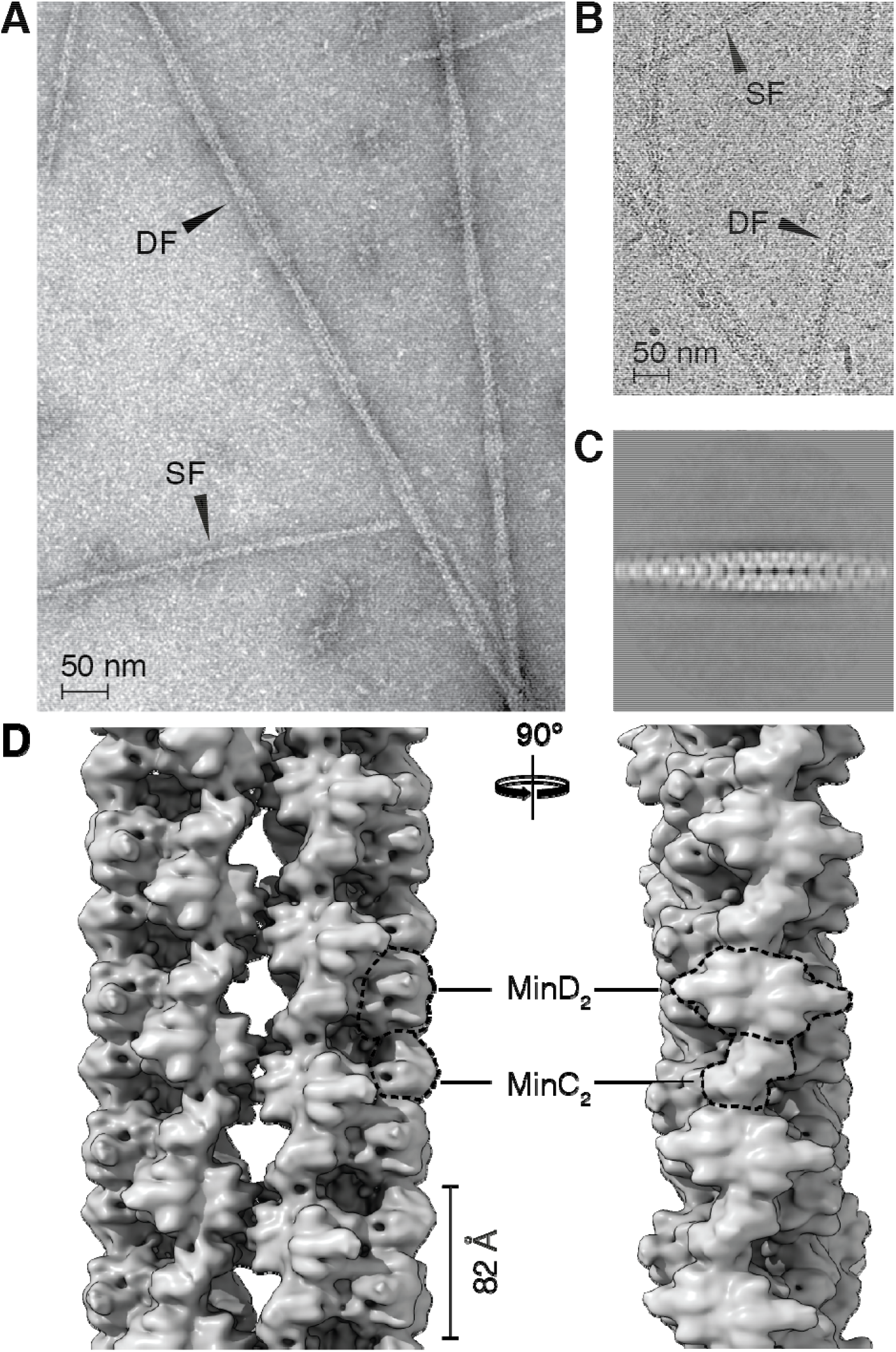
Filaments of *P. aeruginosa* MinCD. (A) Electron micrograph of negatively stained *P. aeruginosa* MinCD filaments, which form upon addition and binding of ATP. The observed polymers of MinCD are a mixture of single (SF) and double (DF) filaments. (B) Single (SF) and double (DF) filaments of MinCD from the same sample, after vitrification, visualised using cryo-EM. (C) Representative average image from reference-free 2D classification and alignment of double filament segments showing the helical nature of the filament. (D) Two orthogonal views of the cryo-EM map resulting from 3D reconstruction of the *P. aeruginosa* MinCD double filament. Two single MinCD filaments wrap around each other in a double helix, but the filaments are helical themselves and made of three MinCD protofilaments each. The protofilaments are built from alternating MinC and MinD dimers, as indicated (dashed lines).

We observed that MinCD filaments were found in the expected mixture of single and double filaments, both in negatively stained as well as vitrified samples observed by cryo-EM. Initially, we focused our attention on the double filaments, which in the collected micrographs appeared to exhibit the helicity required for reconstruction. However, when we carried out reference-free 2D classification and 3D reconstruction of the selected filament segments, it became clear that what appeared to be single filaments making up the two strands of the double helical filament, were in fact themselves helical filaments (Fig. 1C–D). Since these helical, thinner filaments were the dominant species in the polymerised sample, and they also appeared to be more ordered, we focused on them as targets for high-resolution helical reconstruction.

The first round of 2D classification and averaging of MinCD single filament segments showed MinC and MinD dimers (Fig. 2A) arranged in a pattern discernible also in previously published negatively stained micrographs of *E. coli* and *A. aeolicus* MinCD filaments [25]. Close inspection of the 2D class averages and the 3D reconstruction of the double filament revealed that the MinCD single filament is composed of three protofilaments, wrapped around each other to form a very gently twisting triple helix (Fig. 2G). Each of the protofilaments is built from an alternating, copolymeric assembly of MinC_2_ and MinD_2_ dimers. Because of the open-ended (translational) and dimeric nature of MinCD interactions, the filament has two-fold (C_2_) symmetry axes perpendicular to the main filament axis and thus no overall polarity.

**Fig. 2.**
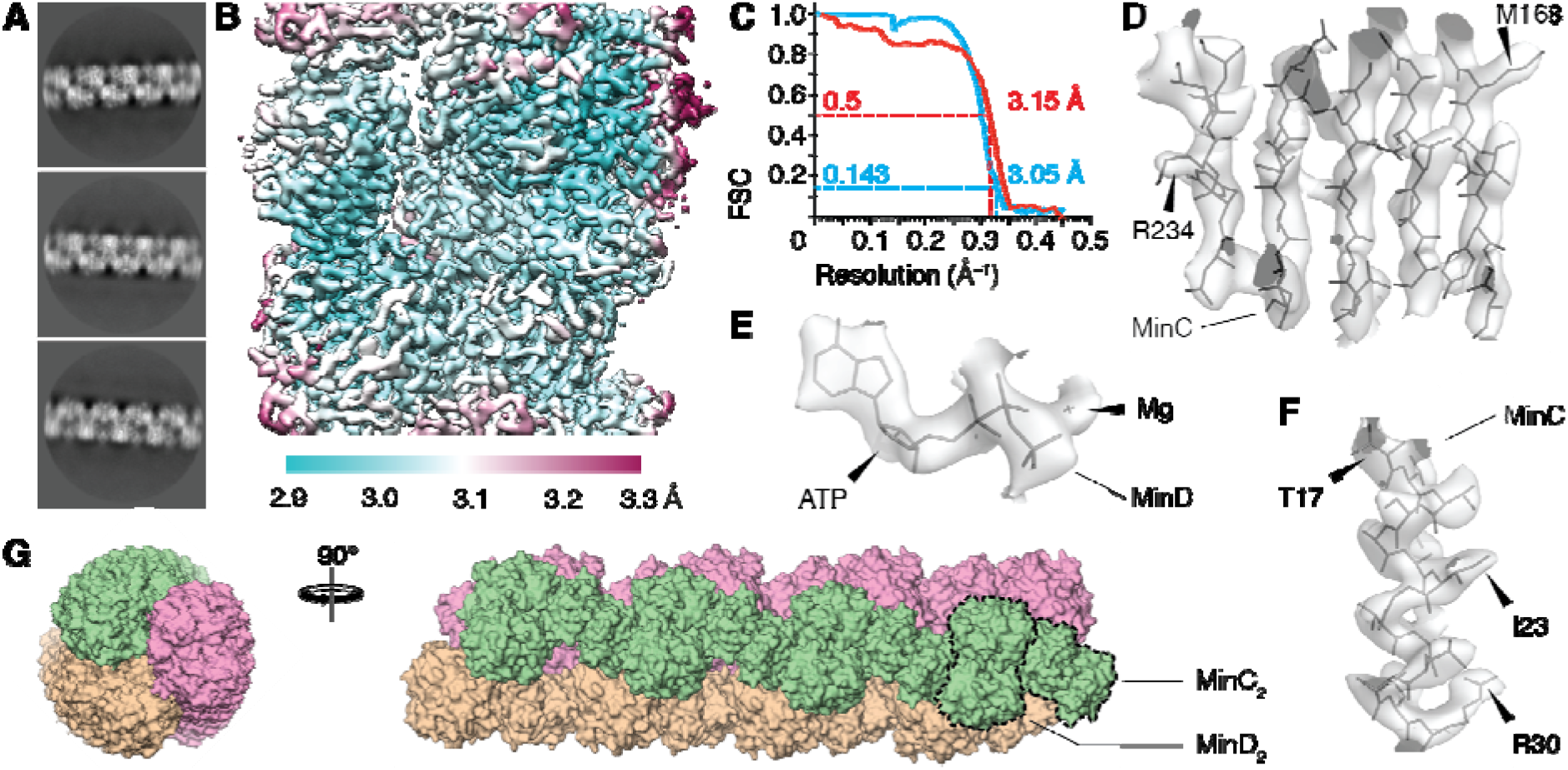
Structure of *P. aeruginosa* MinCD single filament. (A) Representative average images from reference-free 2D classification of single MinCD filament segments. (B) Reconstructed cryo-EM map of the central portion of the MinCD filament, coloured by local resolution. (C) Fourier-shell correlation (FSC) curves between half-datasets (blue, goldstandard) as well as between the map and the model (red). The overall resolution is marked, as judged by the FSC_0.143_ and FSC_0.5_ criteria (dashed lines). (D–F) Selected portions of the reconstructed cryo-EM map with the MinCD atomic model built in: (D) β-sheet, (E) α-helix and (F) the ATP nucleotide with coordinated magnesium. Arrowheads mark the position of arbitrarily selected amino acid residues. (G) Surface representation of the reconstructed MinCD filament model in side and top views. The triple helical nature of the single filament, as well as the repeating MinC and MinD dimers are discernible and indicated in the figure (dashed lines). The three protofilaments are coloured to highlight the arrangement and slight twist of the protofilaments (3.7° per heterotetramer).

3D refinement of the selected helical segments allowed us to reconstruct a cryo-EM map of the MinCD filament (Fig. 2B). The helical parameters of the right-handed MinCD helix were refined simultaneously, yielding values of −25.0 Å (rise) and 116.3° (twist) (Table 1). 116.3° is close to 120°, describing ideal helical symmetry with three protofilaments, and means that the filaments twist gently with right-handedness. The overall resolution of the obtained map was 3.1 Å (Fig. 2C), calculated using the FSC_0.143_ gold-standard criterion, with secondary structural elements and individual amino acid residues resolved to a degree expected from a map at this resolution (Fig. 2D–F). The reconstruction presented here provides high-resolution structural evidence that strongly supports previous proposals that MinCD forms copolymeric filaments composed of alternating protein elements [25,29].

### MinCD filaments are made from a structurally conserved heterotetramer

The resolution and quality of the obtained reconstruction enabled us to build a high-quality atomic model of *P. aeruginosa* MinC and MinD, which was fitted into the cryo-EM map of the MinCD filament and refined. The refinement statistics for the MinCD structure solution are summarised in Table 1, together with data collection parameters. In the final map, almost all of the MinD structure is well resolved, however the last 16 amino acid residues of the C-terminus could not be fitted, due to poorly defined density. For MinC, only the C-terminal dimerisation domain MinC^C^ could be built, accounting for around 40% of the full-length MinC protein. The MinC^N^ domain, thought to be the major site of interaction with FtsZ, is entirely missing in the reconstructed map, most likely due to flexibility (Fig. 3A).

**Fig. 3.**
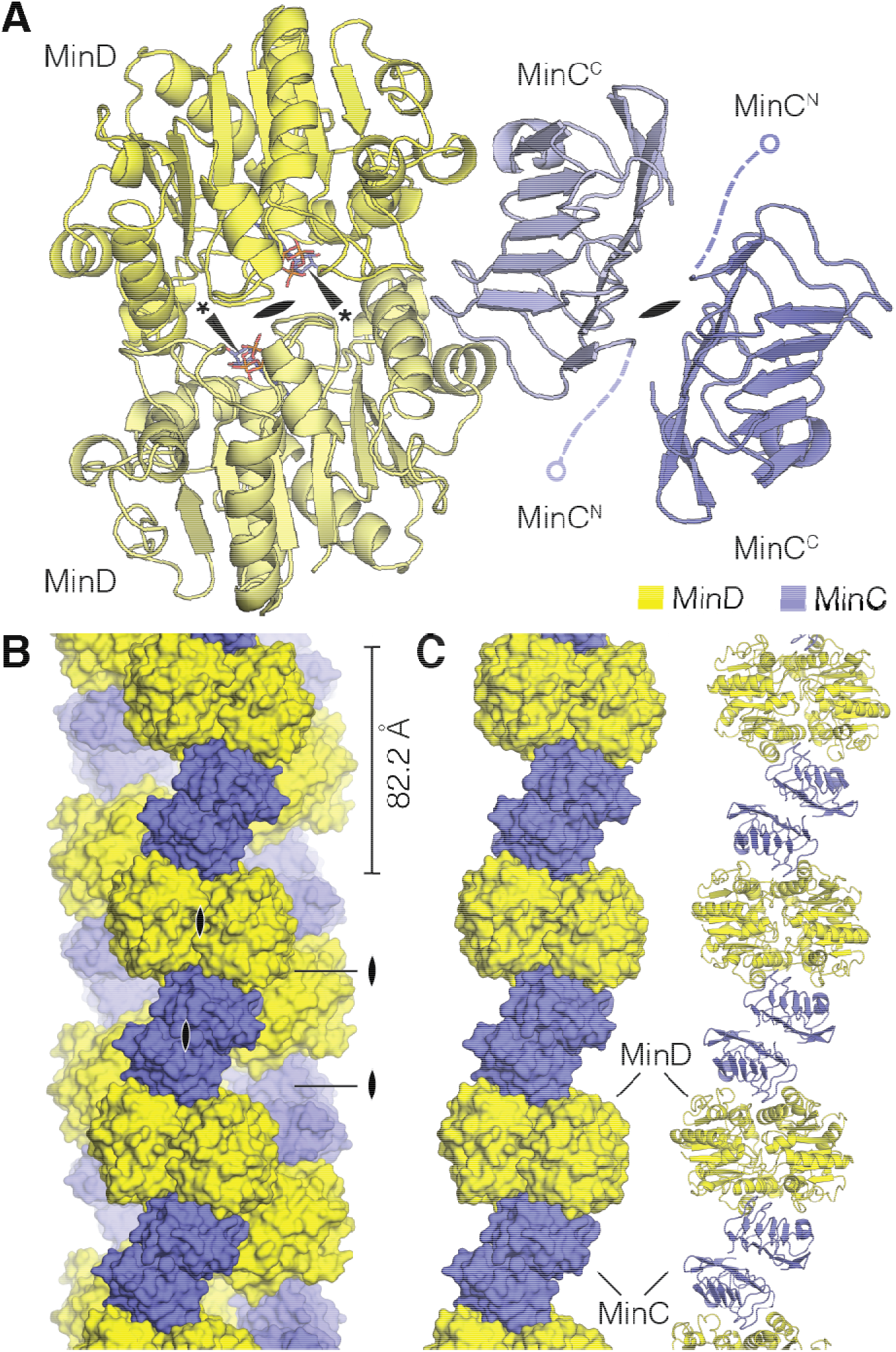
Structure of *P. aeruginosa* MinCD heterotetramer. (A) Atomic model of the MinCD heterotetramer, in cartoon representation, built from and refined against the reconstructed cryo-EM map of the filament. The MinCD tetramer is composed of MinC_2_ and MinD_2_ homodimers, which have been labelled and coloured (yellow and blue, respectively). The positions of the disordered MinC^C^ domains are indicated schematically, as is the position of the 2-fold symmetry axis in each of the dimers (lens symbol). (*) ATP bound to MinD. (B) Surface representation of the atomic filament model coloured according to the scheme from the previous figure panel. The alternating arrangement of MinC and MinD dimers is clear, with multiple 2-fold symmetry axes between any two MinC or MinD monomers (lens symbols). The repeat distance of the tetramer is indicated. (C) One of the three protofilaments from the previous panel, in surface and cartoon representation, showing the arrangement of the tetramers in a single protofilament.

The structure of *P. aeruginosa* MinD shares very strong structural similarity with *E. coli* and *A. aeolicus* homologues, which is not surprising, given respective 74% and 45% sequence identities between the proteins. MinD is a deviant Walker A cytoskeletal ATPase (WACA) and members of this family depend on sandwiched binding of two ATP molecules for dimerisation, and rely on accessory proteins to stimulate their ATPase activity [46]. In our MinD structure, as in the published crystal structures of two homologues [25,47], the catalytic pocket contains non-hydrolysed ATP, added in the experiment to promote dimerisation (Fig. 2F). Without stimulation from MinE, MinD does not readily hydrolyse the nucleotide [19]. Just as in *E. coli* and *A. aeolicus*, the *P. aeruginosa* MinCD filaments are disrupted by MinE, and this is thought to happen in part because ATP hydrolysis causes the MinD dimers to separate, depolymerising the cofilament [25,27].

In our reconstruction, *P. aeruginosa* MinC forms a homodimer via its dimerisation domain MinC^C^, similar to other structures of MinC protein homologues [23,24,48]. The MinC^C^ domain contains a conserved β-helix fold, which has two interaction surfaces: on one side it dimerises with another MinC^C^ and on the other it forms a heterodimer with a MinD monomer (Fig 3A). The MinC^C^ domain is connected to the MinC^N^ domain via a flexible linker, which means that MinC^N^ protrudes out of the filament and is not found in a defined relative orientation with respect to the rest of structure. This explains why, in our map, MinC^N^ could not be averaged into a defined density, although it is present in the protein sample and micrographs.

### Structural features that enable MinCD to form an open-ended polymer

Consistent with previous studies and models, both MinC and MinD are dimeric and form an open-ended polymer in which each protein monomer interacts with its homodimeric partner on one side and the heterodimeric partner on the other. Each of the polymers of alternating MinC_2_ and D_2_ dimers forms a protofilament, and the protofilaments are interacting with each other in such a manner that every MinCD heterodimer interacts with a complete MinCD heterodimer on the neighbouring protofilament (Fig. 3B–C). As mentioned before, the MinCD filament has rotational two-fold *C*_2_ symmetry, which means that two kinds of two-fold symmetry axes are present in each protofilament, but also between any two of the three protofilaments.

The final 16 residues of the MinD C-terminus contain a membrane targeting sequence (MTS): an amphipathic helix with which MinD anchors itself into the membrane upon ATP-binding and dimerisation [17,18]. The MTS of *P. aeruginosa* MinD is part of the unresolved C-terminal region of the protein in our reconstruction. This region exhibits inherent structural flexibility or multiple discrete conformations, which precluded model building. Interestingly, considering filament topology, the MTS of MinD ought to be located in the cavity formed inside the MinCD triple helix (Fig. 4A). We propose that the association of the MTS from MinD monomers in the filament interior is due to non-specific interactions between hydrophobic residues and likely helps to stabilise the three protofilaments in their forming one helical MinCD filament A similar effect has recently been observed in an unrelated bacterial filamentous protein: bactofilin. Bactofilin contains a conserved hydrophobic N-terminal tail, which tethers it to the cell membrane. In the absence of a lipid bilayer, bactofilin assembles into long, double helical filaments, with the N-terminal tails associating in the centre of the filament and stabilising the interaction between the protofilaments [49].

**Fig. 4.**
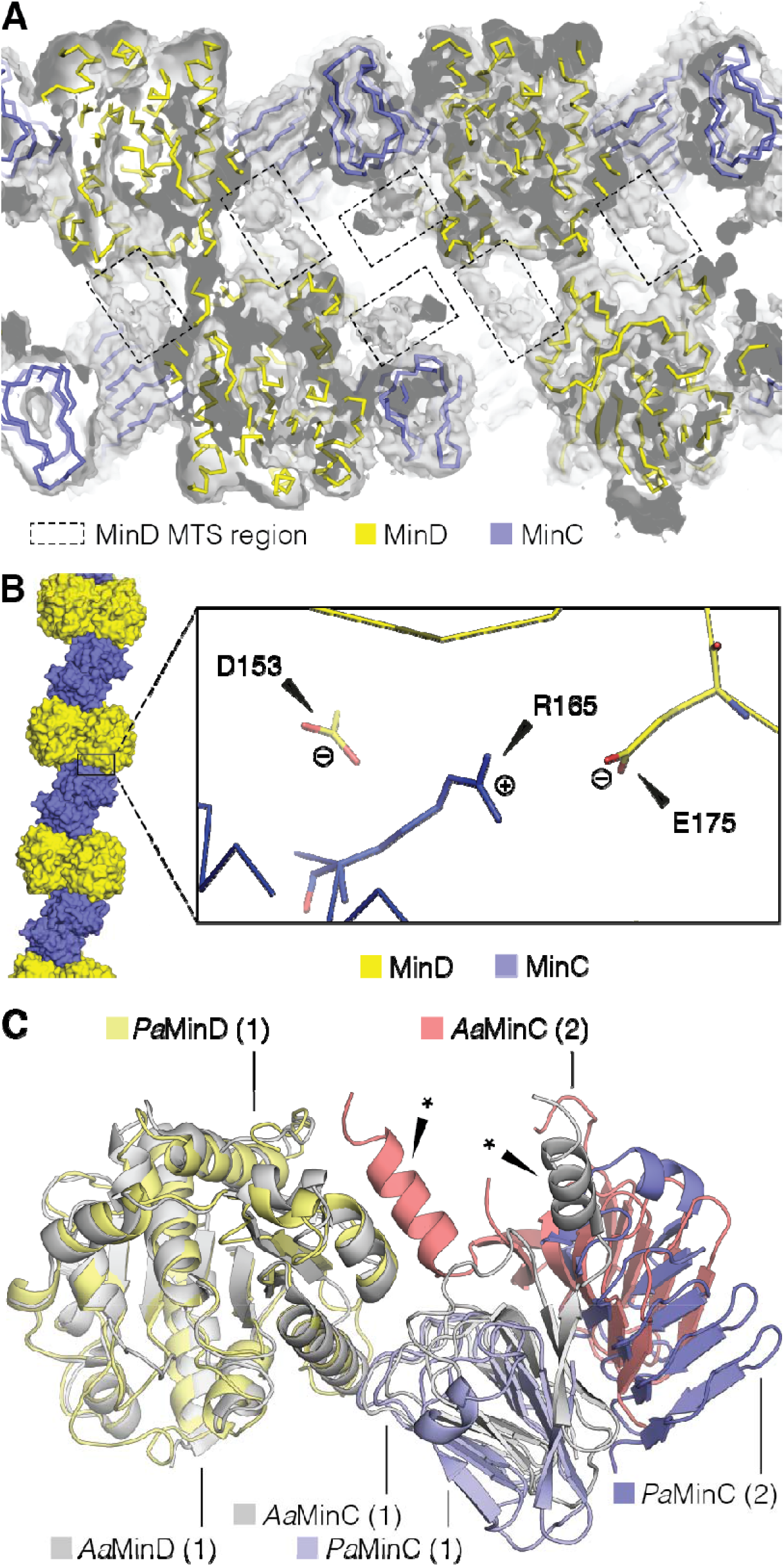
Structural features of the *P. aeruginosa* MinCD filament. (A) Cross-section through the reconstructed cryo-EM map, contoured at σ level 1.0 with the atomic model of MinCD filament shown in ribbon representation. The proteins have been coloured according to the scheme from the previous figure. The regions of poorly defined density (dashed-line boxes) correspond to the C-terminal membrane-binding amphipathic helices (MTS) of MinD that interact non-specifically to bridge between adjacent MinD monomers. (B) Detail of the cryo-EM model around conserved residue R165 of MinD (right) found at the interface of MinC and MinD monomers in the protofilament (left, inset box). This residue is in close proximity to two negatively charged residues of MinC: D153 and E175, with which it might be forming salt bridges. Colouring scheme for MinCD as above. (C) Overlay of MinCD structures from *P. aeruginosa* and *A. aeolicus* (PDB ID: 4V02). The crystal structure of *A. aeolicus* MinCD contains a MinC dimer bound to a MinD monomer so it was aligned with the homologous protein chains from *P. aeruginosa* MinCD tetramer. The proteins are coloured according to the key in the figure. *A. aeolicus* and *P. aeruginosa* proteins show high degree of structural similarity, the main difference being in the presence of a C-terminal α-helix (*) in *A. aeolicus* MinC and providing an additional contact between MinC and MinD.

In the previously reported MinC and MinD co-crystals from *A. aeolicus*, the only other structure of a MinCD heterodimer, the authors identified MinC residue D155 as crucial for MinC and MinD interaction [25]. This aspartate is also conserved in *E. coli* and *P. aeruginosa* MinC at positions D154 and D153, respectively. Mutation of the conserved aspartate into alanine abolished filament formation of the *A. aeolicus*, as well as *E. coli* proteins [25,30]. This is because, as deduced from the crystal structure of *A. aeolicus* MinCD, the residue forms a salt bridge with a conserved arginine of MinD: R94 in *A. aeolicus* and R133 in *E. coli*. In our reconstruction of *P. aeruginosa* MinCD heterodimer, the same salt bridge seems to be present, between D154 of MinC and R165 of MinD. Because of the quality of the map in this region (the external side of the filament), it cannot be excluded that the bridge is also/instead formed with E175 of MinC (Fig. 4B).

In the *A. aeolicus* MinCD heterodimer the heterotypic interaction is also mediated via the C-terminal α-helix of the MinC^C^ domain, which contacts a MinD monomer (Fig. 4C). Sequences equivalent to the *A. aeolicus* C-terminal MinC^C^ helix are not present in most other MinC proteins [23,25,48]. The helix is absent in *P. aeruginosa* MinC, so our conclusion is that the helix is not necessary for filament formation and probably evolved in to add stability to the MinCD heterodimer association.

### MinCD forms straight protofilaments on lipid bilayers

Binding of the MinD dimer to the membrane of the bacterial cell is a crucial feature of the Min system. In its active, ATP-bound state, MinD attaches itself to the lipid bilayer via the C-terminal MTS and in doing so can take MinC dimers as passengers. In this membrane-bound state, the two proteins can also form filaments, as shown by electron cryo-tomography of lipid vesicles decorated with *E. coli* MinCD [25]. This was the first demonstration that MinCD complexes form co-polymeric assemblies not only in solution, but also when bound to the surface of membranes, facilitated by MinD’s MTS.

To test whether *P. aeruginosa* MinC and MinD proteins bind to lipid bilayers we polymerised MinCD filaments in the presence of lipid nanotubes, made from *E. coli* total lipid extract doped with galactosylceramide. Thanks to this addition, lipid nanotubes have a permanent and uniform curvature, forming a rigid and ordered membrane-like support for membrane-binding proteins, which can facilitate electron microscope imaging [50–52]. Lipid nanotubes have already proven useful in the study of cell membrane assembly of another protein which forms prokaryotic cytoskeletons, the bacterial actin MreB [53].

Cryo-EM images of *P. aeruginosa* MinCD filaments polymerised with lipid nanotubes clearly show that the filaments assemble on lipid bilayers (Fig. 5A–B). The tubes are evenly coated with MinCD filaments, and the absence of free filaments suggests that the interaction of filaments with the lipid environment is more stable than the self-interactions, which generate the three-stranded filaments in solution as resolved by cryo-EM. Due to its geometry, the edge of the nanotube is where the presence of the filaments is most pronounced, with least overlapping of filaments, allowing us to carry out reference-free 2D averaging of the nanotube surface with MinCD bound to it.

**Fig. 5.**
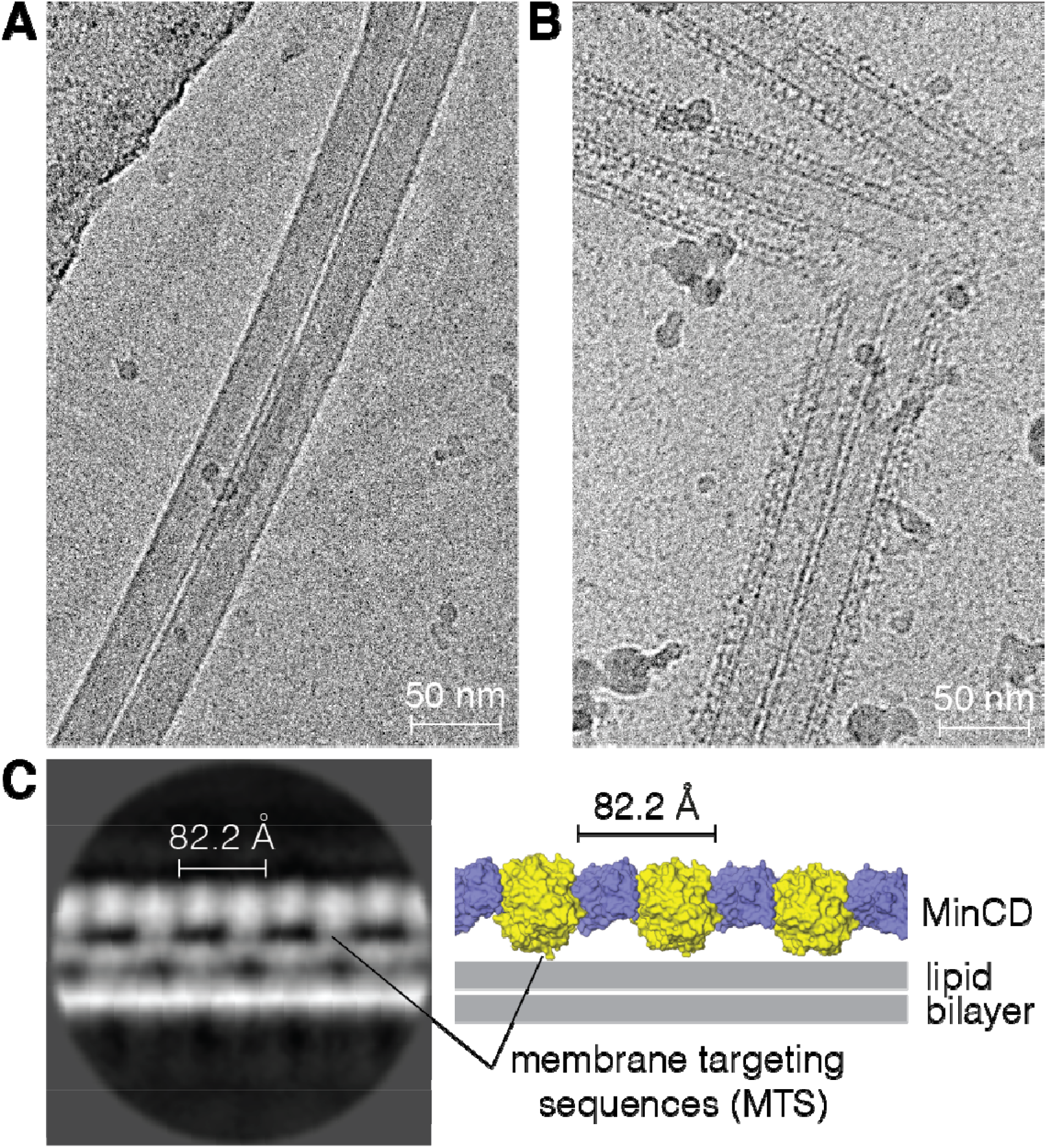
Imaging of *P. aeruginosa* MinCD filaments binding to lipid nanotubes. (A) Micrograph showing two undecorated (protein-free) lipid nanotubes imaged after vitrification using cryo-EM. (B) Similar image, with added MinCD and ATP. An ordered layer of protein decorating the surface of lipid nanotubes is clearly visible compared with the previous panel. (C) A representative average image from reference-free 2D classification of the lipid nanotube surface segments (left). The MinCD filament is bound to the lipid bilayer on the outer surface of the tube, also represented schematically with the MinCD protofilament atomic model derived here (right). The characteristic alternating pattern of MinC and MinD dimers with an 82-Å repeat (cf. Fig. 3B) is discernible in the 2D average image. The approximate positions of the membrane targeting sequences (MTS) of the MinD dimers are indicated. The distortions seen in both leaflets of the lipid bilayer are likely due to the interference from MinCD protofilaments bound to the tube surface above and below the plane of the reconstruction.

The alternating co-polymeric assembly of MinCD dimers on the lipid surface is apparent in the 2D class averages of the micrographs (Fig. 5C). MinD dimers are tethering the filaments to the membrane, and two attachment points, each from an MTS from a MinD monomer, are also discernible. The MinC dimer does not contact the membrane, but links the MinD dimers together, just as suggested by our reconstruction of unsupported MinCD filaments.

MinCD assembled on the surface of a lipid bilayer does not form a triple helix, as in solution, but rather appears as single MinCD protofilaments. Instead of a helix with protofilaments facing each other on the side of the MTS of MinD, protofilaments arrange into straight assemblies anchored by the MTS on the surface of the lipid nanotube. The slight helical twist of the triple helix (3.7° per heterotetramer, cf. Fig. 2G) is either reduced or absent, as the membrane-bound filaments seem to be perfectly parallel to the long axis of the tube (Fig. 5C). Some degree of flexibility in MinCD subunit interaction necessary for such adjustment may not be without precedent, as *A. aeolicus* MinCD het erodimers in the crystal structure are bent, but have been reported to also form straight protofilaments on lipid bilayers [25].

## Discussion

Our previous hybrid model for *E. coli* and *A. aeolicus* MinCD filaments [25] had the two homodimeric C_2_ axes lying in a single plane, both perpendicular to the tangent of the curved filament implied by the crystal structures. We suggested that unbending of the curved polymer, to bring these axes parallel, would therefore expose several MinD MTS in a colinear arrangement, allowing efficient and avid binding to membrane – explaining the observation of straight filaments on lipid vesicles *in vitro*. The work here demonstrates that these predictions were correct and generalise to another MinCD pair. Firstly, the structure of MinCD cofilaments from *P. aeruginosa* shows that MinCD can form straight, untwisted, filaments in the manner expected. Further, visualisation of MinCD bound to lipid nanotubes demonstrates that these straight filaments indeed do use the linear array of MinD MTS sequences exposed along one surface to assemble efficiently on membranes.

It must be emphasised that the coplanar orientation of the homodimer C_2_ axes we have now observed in several MinCD pairs, and which is essential for avid cofilament membrane binding, is not an inevitable geometric consequence of the interaction of two C_2_-symmetric dimers, instead it is a rather special arrangement which can only be satisfied by a narrow range of heteromeric interaction modes. Notwithstanding the compelling evidence of Park *et al*. [30] that MinCD copolymers are not essential for Min function in *E. coli* cell division inhibition, it remains challenging to explain why this special relative arrangement of MinCD dimers would persist, especially as open and purely translational symmetries are usually avoided in cells unless polymerisation is selected for during evolution. We believe that it remains a distinct possibility that membrane-bound MinCD cofilaments do in fact play a role in cells, and that they could therefore be part of an expanding class of non- (or hardly-) twisting membrane-binding protein polymers found in bacterial cells, including filaments formed by MreB, FtsA, bactofilin, SepF, and possibly DivIVA [49,54–57].

## Acknowledgements

We thank the staff of the Electron Bio-Imaging Centre (eBIC) at Diamond Light Source, Didcot, United Kingdom, and the staff of the MRC-LMB Cryo-EM facility: Shaoxia Chen, Giuseppe Cannone and Joanna Brown, for their assistance with data collection. We would also like to thank Jake Grimmett and Toby Darling, from the MRC-LMB Scientific Computing Facility. This work was funded by the Medical Research Council (U105184326 to JL) and the Wellcome Trust (202754/Z/16/Z to JL).

## Author contributions

ASH did all biochemistry, ASH and JW did electron microscopy, JL, ASH and JW did image analysis and reconstruction, JL did model building and refinement. ASH wrote the manuscript. All of the authors approve this submission and declare that they have no competing interests.

## Abbreviations

Cryo-EM: electron cryo-microscopy
FSC: Fourier shell correlation
IPTG: isopropyl β-d-1-thiogalactopyranoside
WACA: Walker A cytoskeletal ATPase
TEM: transmission electron microscope
MTS: membrane targeting sequence

